# Emergent organization of multiple visuotopic maps without a feature hierarchy

**DOI:** 10.1101/2021.01.05.425426

**Authors:** Talia Konkle

**Affiliations:** Harvard University

## Abstract

The primate visual system is comprised of multiple visual areas. Despite their foundational relevance, there are no normative accounts for why there are multiple areas nor why they have their signature “mirrored map” topography. Here I show that the stereotyped cortical organization of multiple mirrored areas naturally emerges in simulated cortex, in which self-organizing processes are used to map a multi-scale representation of visual space smoothly onto a two-dimensional cortical sheet. Predominant accounts of multiple areas emphasize hierarchical processing, where each area extends and elaborates on the previous areas’ representation. Here, no explicit hierarchical relationships were required to manifest this multi-areal organization, suggesting that feature hierarchies may be the derived rather than the driving force of this organization. This modeling work thus provides a simple computational explanation for the hallmark features of early visual topography, and the presence of multiple areas, as emergent from a single functional goal — to smoothly represent the visual field at multiple spatial scales.

**One Sentence Summary:** This work presents a formal model of simulated cortex with multiple visual areas, where purely spatial relationships underlie the large-scale motifs of visual cortex.

---

A foundational empirical observation of the visual system’s earliest cortical processing stages is that neurons that are nearby on the cortical sheet also respond to light hitting the retina from nearby parts of visual space, forming multiple “visuotopic” maps along the cortex (*1*). Primary, secondary, and tertiary visual areas V1, V2, and V3, each have their own representation of the visual hemifield (*2*). The hierarchical relationship among these areas is perhaps their most well-known property, conceived of as a cascading series of distinct and separate processing stages, each with their own representational goal (e.g. *3,4*). However, this conceptualization of visual areas does not account for their systematic, intricate, and integrated visual field topography along the cortical sheet (**Fig. S1)**. Inspecting this unified topography have led some to proposed that there may be a single common functional role which unifies these areas (see *6,7*; see also *8,9*). However, to date there are no formal models that articulate what the joint functional role of this map complex might be, nor any that predict the large-scale motifs of the multi-areal mirrored map organization of early visual cortex.

Here I take a representational mapping approach, drawing on the same methods that have been used to predict the fine-scale orientation pinwheels embedded in retinotopic V1 (*10*), and the large-scale organization of body part and movement tuning in the motor-premotor complex (*11*). These approaches leveraged a self-organizing map algorithm (SOM) developed by Kohonen (1989; *12*). This computational framework takes as input a high-dimensional representational space, and outputs a tuned map, in which nearby units in the map project to nearby points in the representational space (learning a smooth 2-dimensional embedding to capture the structure of the input space). Theoretically, a smooth mapping of information onto the cortex may be desirable because it enables local computations in the parameter space to be performed by local circuits in the cortex (e.g. visuotopic maps would enable center-surround computations to be carried out with a local, repeating connectivity motif). Further, the mapping solutions obtained with the self-organizing map algorithm create an approximate solution to the NP-hard problem of arranging connected units in a spatial grid to minimize overall wiring (*10*; *13*), but accomplish this with simple, biologically inspired, activity-dependent learning rules (*12*).

Cast in this computational framework, the key question becomes: what is the representational space that, when smoothly mapped, gives rise to the multiple mirrored maps of the early visual cortex? Here I focus on the structure of a multi-scale representation of visual space, which has been foundational for computational approaches to visual image analysis, given the structure of natural images (e.g. *14*). To anticipate, I show that a simple multi-scale representation of visual space, when smoothly mapped, can account for the major large-scale motifs of visual cortex organization. The simulated cortical sheet shows emergent “areas,” defined based on mirrored alternations of polar angle meridians, despite the fact that no explicit hierarchical feature relationships are specified. These emergent “areas” also naturally show well characterized receptive field size properties, which increase with area and depend on eccentricity. Thus, this modeling work provides the first account of the supra-areal organization of early visual cortex, and proposes that multi-scale spatial relationships alone are sufficient to account for the arealization of cortex, without an explicit feature hierarchy.

The key simulation results are shown in **Figure 1**. For the main simulation, an implicit multi-scale filter bank was defined as the input (**Fig. 1a**). Specifically, first the visual field was discretized into a large number of locations, where each point in the visual field is a separate dimension—a key contrast with prior work which defined visual space more explicitly with two dimensions (e.g. *10, 15*). With this parameterization, any potential input can be described as a vector in this space, based on its activations at each visual field location. Next, a series of 2-dimensional Gaussian input samples were created at each visual field location, at 8 different scales. For example, a single input sample is an 820-dimensional vector with high values on some locations and lower values on others, with 6,560 samples in total (see **Supplementary Methods)** A visualization of this high-dimensional input space along its first three principal components is shown in **Figure 1b**, revealing a compact and structured representational geometry evident in the higher-dimensional space of possible activation patterns. The goal of the SOM algorithm is to find a smooth 2-dimensional embedding that spans this input space.

**Fig 1.**
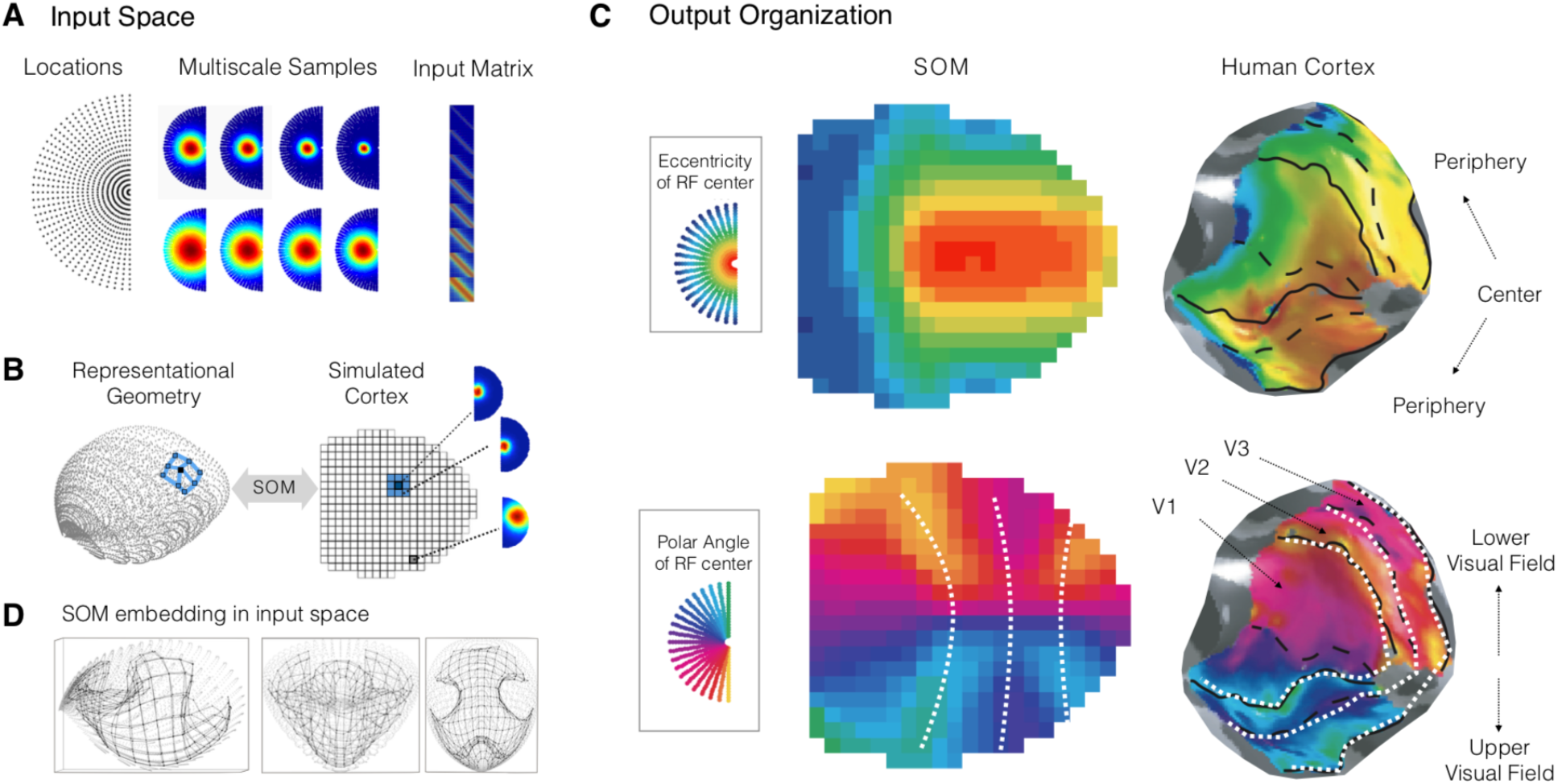
Emergent multiple mirrored map organization. (A) 820 dimensions were used to discretize locations in the visual hemifield, and input samples were created as 2-dimensional Gaussians centered at each location in the visual hemifield, at 8 different scales. One sample is shown at each scale for illustrative purposes. The resulting input space of 6560 samples is visualized in matrix form; and in (B) it is plotted along its top three principal components. The goal of the SOM algorithm is to project a 2-dimensional grid of units into this input space (e.g. blue unit), where nearby units in the SOM are tuned nearby in the input space (light blue units). (C) The resulting organization of receptive field tuning in the simulated cortex (or SOM) is shown. The receptive field center of each map unit is colored by eccentricity (upper row) and polar angle (lower row), with different color scheme shown in the legends (gray boxes). A depiction of the human areal organization is adjacent, adapted from (*39*). Alternations along the horizontal and vertical meridians are schematized with dashed white lines. (D). Visualization of the SOM projected into the input space. Each gray dot is an input sample, and each black dot is a map unit, where adjacent map units are connected with black lines. The same projection is shown from different viewpoints to visualize how the 2-dimensional map manifold fits in the input space.

Submitting this input space to the SOM algorithm yields a tuned “map” (or simulated cortex), which is comprised of units arranged in a grid, each with a set of learned weights (or tuning curves) tuned in 820-dimesional space. The modeling procedure places no constraints on the resulting tuning of the map units—thus, any unit could have any possible spatial tuning, e.g. small or large receptive fields, or non-Gaussian tuning like multi-lobular receptive fields or eccentricity rings. However, inspecting the implicit tuning of each map unit revealed that the learned receptive fields were well characterized by 2-dimensional Gaussians (**Fig S2**). The tuning of each unit in terms of its eccentricity, polar angle, and size in the simulated cortex was then fit with a 2-dimensional Gaussian.

The resulting visuotopic organization of the simulated cortex is shown in **Figure 1C**, adjacent to human visual field maps of the V1-V2-V3 areal complex. The hallmark large-scale motifs of these early retinotopic areas spontaneously emerge. That is, there is a large-scale organization by eccentricity, with a center-to-periphery organization running along one axis of the map (**Fig 1C**, top row). Second, there is a major division between the upper and lower visual field along the orthogonal axis of the map (**Fig 1C**, bottom row, tuning by polar angle). And critically, there are emergent alternations of horizontal and vertical meridian representations, which is the functional signature of areal boundaries (**Fig. 1C**, dashed white lines, *(1)*). Thus, the first key result is that smoothly mapping a multi-scale filter bank is sufficient to give rise to the multiple mirrored map motif of the early visual cortex in humans (and many other primates *(8)*).

Based on these alternations, I defined “areas” in the simulated cortex, analogous to how visual areas V1, V2, and V3 are functionally defined in human and other primate visual systems following the meridian reversals. Two further characteristic signatures of early visual cortex organization were evident (**Fig. 2**). First, the receptive field sizes of map units in the first area were the smallest, and increased with successive areas. Second, in each area, receptive field sizes also increase systematically as a function of eccentricity (**Fig. S11**). Thus the systematic relationships between scale, eccentricity, and areas also spontaneously emerged in this simulated cortex. A somewhat hidden but important implication is that there is not a one-to-one mapping between areas and spatial scale (e.g. in this example there are more scales present in the input space than there are emergent areas, and each area contains a systematic range of scale space).

**Fig 2.**
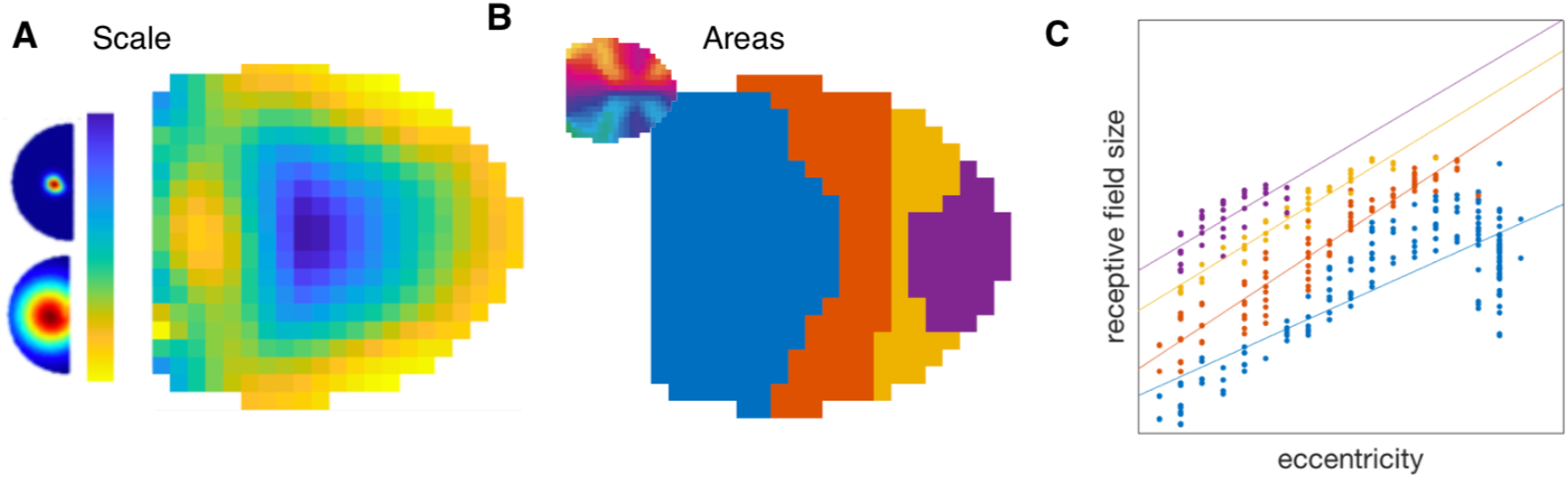
Relationship between emergent areas and receptive field size. (A) The receptive field size of each map unit is shown, where units with smaller receptive fields are located at the center of the map. (B). Emergent areas are visualized, defined based on the polar angle alternations (inset). (C) Plot of the relationship between eccentricity and receptive field size, colored by emergent area. Each dot is a map unit, plotted by fitted eccentricity and scale parameters. Lines show least squared fits of these relationships, separately for map units from different areas.

This emergent multi-areal organization is surprising because the multi-scale filter bank does not have any explicit, progressive feature hierarchy. For example, the input space does not specify any feature tuning relationships (e.g. from simpler orientation features to more complex conjunctive features). So, why does this simulated cortex have “areas” (*3, 16*)? This modeling work indicates that the mirrored alternations are an emergent consequence of trying to simultaneously smoothly map multiple spatial scales. In the final map, the smallest spatial scale ends up at the center, with increasingly larger scales tiling around it (**Fig. 2A**). In this way spatial scale and eccentricity are the related factors that determine the largest scale organization, with polar angle representations mirroring as the 2-dimensional embedding smoothly weaves through this input space (**Fig. 1D**).

A number of model variations were conducted to further probe the link between multiple scales and emergent areas, revealing a range of more simple or complex topographies. While the exact shape of the map (e.g. semi-elliptical, rectangular) had little effect on the resulting organization (**Fig. S3**), both the size of the map (i.e. number of units or amount of cortical real-estate), and specification of the multi-scale filter bank (e.g. how much scale space, how densely sampled, etc.) interact to determine the final map organization (**Fig. S4, Fig. S5**). For example, the same multi-scale input space, when given more cortical territory, will often show more “areas”, with variability in their sizes (*17, 9*). Some combinations do not yield clear areas but instead have an interdigitated retinotopy, especially when the range of scale space sampled is relatively small, and the amount of cortical real-estate is relatively large. Interestingly, in primate visual systems, even though the large-scale organizational motifs are generally consistent, there is substantial individual variability in the relative size of areas (e.g. V1 varies 3x in humans (*18)*; and the V1-V2 ratio varies across primate species *(19)*), as well as finer-grained retinotopic organizations and deviations (e.g. *20*-*22*). This computational framework may thus have further explanatory power, as it enables predictions about differences in areal sizes and arrangements, related to hypothesized differences in allocated cortical area and spatial scale sensitivity.

Model variations also show that, across nearly all specifications of the multiple spatial scale input space, the major motifs of upper-vs-lower and central-vs-peripheral organizations were present. The reason seems to be that these two dimensions are deeply related to the structure of the multi-scale representational space itself (**Fig. 1B**), rather than the SOM algorithm. For example, these motifs are evident in projections of the same input space by other dimensionality reduction techniques like PCA, t-SNE and multi-dimensional scaling (**Fig. S6, Fig. S7**). However, the polar angle alternations, which are critical for how we functionally-delineate areas, are only evident with the self-organizing map algorithm. One insight into this difference is that the SOM algorithm operates in a different “direction” than the others (see *10*). PCA, t-SNE and MDS all map the input space into 2-dimensional coordinates; SOMs map two-dimensional coordinates into the input space (**Fig. 1D**). In other words, SOMs assume a regularly, locally connected cortical surface and learn an embedding in the input space, while the other methods do not have any explicit cortical manifold in their formulation.

These modeling results primarily offer a normative perspective on why the visual cortical areas take the form they take (e.g. akin to *11*). That is, these results demonstrate that a multiple-mirrored map organization can emerge from minimally assuming the representational goal of a multi-scale coding of the visual field, with a spatial constraint of keeping similar tuning nearby on the cortex. As such, this work implies that quantifying and visualizing receptive field size along the cortical mantle may be an important source of information into the relationship among areas of the visual system, especially in species with different visuotopic motifs (e.g. *23,24*). Further, this work may provide insights into the auditory domain, where there are also mirrored maps of tonotopy (*25*).

It is an open question whether this modeling framework can also be informative for understanding the development of visual areas at a more mechanistic level. Indeed, Kohonen’s self-organizing map algorithm was inspired by biology, and assumes a simple, locally connected cortex, where activity-dependent mechanisms guide the map unit tuning to reflect the statistics of the input. The first stage of training is an initialization as a loose retinotopic hemifield related to the major plane through the multi-scale input space (**Fig. S8**)—in the model, this step is critical for yielding consistent large-scale organizations, without it the emergent organization is highly variable across modeling runs (see **Fig. S9**). In the second training stage, map units are fine-tuned by experiencing multi-scale inputs, each unit competing with others to represent different inputs; **Fig. S10, Movie S1** for a video of how the “areas” develop in the simulated map). Interpreted through a biological lens, the first model step is akin to a cortical targeting stage where gradients of morphogens guide retinotopically-organized thalamic inputs to appropriate parts of early visual cortex (*26*). The second model step is similar to subsequent retinal waves, which drive activations in the cortex to sculpt the resulting retinotopy (*27*). Thus, by identifying the task of learning a multi-scale representation, this work has potential to make contact with concurrent accounts of visual cortex and arealization that model the developmental processes at these more mechanistic levels of abstraction (e.g. by considering axon growth (*28*), and the structure and stages of retinal waves over different types of retinal ganglion cells (*29*)).

Overall, these modeling results invite revisiting the construct of a visual cortical area as a distinct functional module, and suggest there may be additional value in considering these areas together as a joint functional unit (*6-8, 30*). Conceived of as a whole, this cortex makes information at different spatial scales *simultaneously accessible*, for example, enabling later object-selective areas to access multiple-scales at the same time by connecting jointly with V1 and V2 and V3. Importantly, this view certainly does not counter the idea that neurons in these areas have different feature tuning and participate in a functional hierarchy—they clearly do (*3*). Instead, these modeling results offer a complementary account of arealization, one where areas can emerge based on spatial relationships alone; as such this account may help provide insight into the developmental puzzle for how later hierarchical areas could emerge before earlier areas have refined their feature tuning (*31-33*).

Broadly, this work highlights the importance of representational topography—of characterizing how the tuning of different neurons changes over the cortical mantle, as a critical empirical source of insights into cortical function. The brain does not start out with all neurons connected to all other neurons, where all large-scale brain organizations are possible. Instead, there are a few long-range routes through an otherwise predominantly local set of connections (*34*). Thus what kind of information comes to be represented where, at a macro scale, becomes hugely important for channeling information between sensory and motor systems, (e.g. cortex adjacent to central visual field representations take on more sensitivity to curved features evident in small manipulable objects and link to parietal visual-motor circuits, while the far visual periphery reflects more rectilinear featural tuning evident in big objects and surfaces and connects to navigation-related networks along the medial surface, *31, 35, 36, 37*). As such, understanding the spatial mapping of information along the cortex may provide direct insight into the functional divisions that evolved to support ecological behavior (*38*).

## Acknowledgments

I thank R. Buckner, R. Born, M. Livingstone, D. Leopold, T. R. Candy, K. Miller, R. Saxe, A. Oliva, G. Alvarez, and one anonymous pre-peer reviewer for their thoughtful comments.

## Author contributions

T.K. conceptualization, methodology, software, writing.

## Data and materials availability

Code required to reproduce the main results will be available on git hub following publication.

## Supplementary Materials

Materials and Methods

Figures S1-S12

Movie S1

## Supplementary Materials for

### This PDF file includes

Materials and Methods

Figs. S1 to S12

Caption for Movie S1

### Other Supplementary Materials for this manuscript include the following

Movie S1

Code Base

## Materials and Methods

### SOM Algorithm Overview

Self-Organizing Maps (Kohonen, 1989; Kohonen 2001) were developed as a dimensionality reduction technique, inspired by biological learning mechanisms. This algorithm takes as input a multi-dimensional space as well as parameters describing the 2-dimensional map surface (i.e. its size, shape, and neighborhood definitions), and the learning process (i.e. how many iterations for training). This algorithm outputs a tuned map, that is the map units are projected into the input space and then their tuning is adjusted so as to capture the structure of the input data as smoothly as possible. In particular, the map units learn their tuning by competing to represent the input samples, and then shifting their tuning towards the best matching input samples and shifting the tuning of their neighbors in the same direction. Because of this procedure, the map tuning is sensitive to the distribution of the data (e.g. more map units become tuned to parts of space that are overrepresented by input samples, and fewer map units to parts of the input space with few input samples).

### Modeling Procedure

In this section I describe the specific method choices used to create the early visual cortex map depicted in the main manuscript. The codebase was written in MatLab, and was developed starting from the SOM Toolbox for MatLab 5 created by Juha Vesanto, Johan Himberg, Esa Alhoniemi, and Hua Parhankangas. http://www.cis.hut.fi/projects/somtoolbox/.

#### Discretize the visual hemifield

To sample the visual hemifield with discrete locations, I used a polar sampling scheme, specifying r radii and n eccentricity steps. In the main example in the manuscript, the visual field was sampled with 41 radii and eccentricities from 1 to 20 in 1 degree steps, yielding 820 visual field locations.

#### Create multi-scale input samples

The next step is to specify input samples over these visual field locations. In the main example, Gaussian activation patterns were created at 8 different scales [2 through 10 degrees in steps of 1], centered at each visual field location. These samples were normalized to have a value of 1 at the center location of the Gaussian and fall off to zero at the most distant visual field location. The aim of this step is to generate a feature matrix, with dimensions *n x d* where n is the number of input samples and d is the number of discrete visual field locations. In our specific example, with 8 scales of inputs centered at each of 820 locations there are 6560 total input samples (8 * 820 = 6560). Thus, in the main example, the feature matrix has dimensionality 6560 x 820.

It is important to note that all visual degree measures are actually relative. For example, in the SOM described in the main manuscript, the hemifield is specified with a range of 1 to 20 degrees with steps of 1 degree, and with input samples specified with widths of 2 degrees to 10 degrees by steps of 2 degrees. This set of samples is isomorphic to a hemifield specified from 1 to 40 degrees in steps of 2 degrees and a scale space sampled from 4 to 20 degrees in steps of 4 degrees. In other words, these input numbers should be interpreted as relative rather than absolute degrees.

To facilitate a more geometric intuition about the structure of this space, **Figure S7** visualizes this input space along its first 3 principal-components. PC1 = 68.8% variance, PC2 = 11.6% variance, PC3 = 8.4% variance. Total variance accounted for by first 3 principal components = 88.8%.

#### Specify the size and shape of the simulated cortex

The next step is to specify the map topology, namely the size and shape of the simulated cortical sheet. The number of units used in the main manuscript was set using the default heuristic in the SOM toolbox:

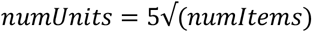

These units are given a location in a 2d grid, specified as map coordinates. The map aspect ratio is calculated based on the ratio of the eigenvalues of the first two principal components. For the SOM reported in the main manuscript, the units were arranged in a partial ellipse, where the eccentricity of the ellipse (i.e. how nearly circular it is) was set as the ratio of the first two eigenvectors, and 75% of the long axis of the map was retained. This yielded 340 map units, filling most of a 20 x 21 grid. This shape was set for aesthetic purposes, and did not have a dramatic impact on the emergent organization (**Fig. S3**).

#### Tune the simulated cortex

During map training, the goal is to establish tuning weights for each map unit, so that each unit is tuned somewhere in the input feature space. The SOM algorithm helps ensure that nearby map units are tuned to nearby points in the parameter space. To train the map there is a two-stage process. First the tuning of each map unit is initialized in the input space. Next, the samples are “shown” to the map units, which adjust their tuning, and their neighbors’ tuning, towards the input samples. The output of this self-organizing map process is the tuning curve of each map unit (*numMapUnits x numLocations*, in our example, 340 x 820). In the code, this output is referred to as the codebook. There were no major innovations or changes made to the underlying algorithm implemented in the SOM toolbox from 2000.

#### Initialize the map

To initialize the map in our example, each map unit is assigned an 820-D vector for its initial tuning. Specifically, the map (or “simulated cortex”) is initialized as a plane along the first two principal components (**Fig. S8a**). In other words, to put an interpretation on this step, the initialization process establishes for each map unit how sensitive it is to different parts of the 820 visual field locations (**Fig. S8b**). This initialization option smoothly initializes the map units to be evenly spaced spanning across the plane formed by the two major axes of the input space. In effect, this initialization helps to situate the simulated cortex approximately to the input space that it will be further tuned to in the next stage of training.

One interesting observation about this initialization step is that a plane along the first two principle components of the input space is effectively a simple visual field map (**Fig. S8c**). That is, the units at the top edge of the map have sensitivity to visual field locations at the center of the hemifield, while units at the bottom edge of the map have sensitivity to visual field locations at the periphery (**Fig. S8c**, left subplot). Similarly, units at the left of the map have lower-visual field tuning, and units at the right of the map have upper visual field tuning (**Fig. S8c**, middle subplot). The variation in scale is less pronounced after this initialization stage, with most units having moderately sized receptive fields (**Fig. S8c**, right subplot), with larger and less 2-dimensional receptive fields at the base of the map (**Fig. S8b**). There is also a hint of a correlation between eccentricity and receptive field size, with slightly larger receptive fields in the units with more peripheral tuning.

The alternate initialization option is to randomly initialize each map unit tuning somewhere in the feature space, rather than initializing each one along the first two principal components. Random initialization leads to fractured large-scale organization (**Fig. S9**).

#### Fine-tune the map

After initialization, the next step is to modify the tuning functions of each map unit so that it is a tighter fit to the data manifold. In other words, the training procedure adjusts each map unit’s tuning to be nearer to the data samples, rather than tuning in a part of the input space that is possible but not prominent in the input, like a checkerboard pattern over visual field locations. Describing the algorithm intuitively, for each data sample in the input space, the best matching map unit is found (i.e. that is, the unit whose current tuning is closest to the particular data sample). Then, for each map unit that is a best-matching-unit, the 820-dimentional vector is adjusted in the direction of the data samples it best matches, and the map units in a neighborhood around each unit are similarly adjusted. In the *batch* method of training, all input samples are matched to the map units, and all map units are updated in one step.

This algorithm is implemented more efficiently in the SOM toolbox (see the technical documentation of the original SOM toolbox for more information, Alhoniemi et al., 2000). Specifically, the input samples are first partitioned according to Voronoi regions of the map tuning (at its current state). Next, the sum of the input samples in each Voronoi set is calculated for each map unit:

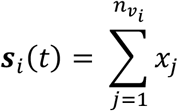

where *n*_*vi*_ is the number of input samples *x* in the Voronoi set of the map unit *i*. Then, the new map unit tuning weights are calculated for each map unit at the next time step as:

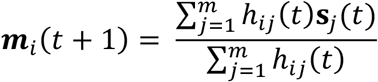

where *m* is the new tuning of the *i* th map unit, at iteration time step *t*, for *m* total map units, and where *h*_*ij*_(*t*) is the neighborhood kernel around the *i* th map unit.

The process is repeated iteratively until the pre-specified number of iterations has been reached. At each iteration, a measure of map quality can be computed, which quantifies how far the map units’ tuning functions are from the input feature space. Tracking this value over iterations indicates whether the map tuning has stabilized. The map quality (qe) for the SOM reported in the main manuscript is plotted as a function of training iteration in **Fig. S11**.

There are several parameter choices related to the *batch* training method, that specify the size and shape of a map unit’s neighborhood (*h*_*ij*_), and how that neighborhood should change over iterations, (e.g. linearly, step function, or not at all), and how many training iterations should be done. In the present work, the neighborhood radius was fixed at 1 for the entire training procedure and was not changed. The number of iterations was selected to be large enough so that the map had stabilized its configuration. The impact of different neighborhood functions and learning rates was not explored in the current work.

#### Visualize the organization of the tuned cortex

At the end of training, each map unit’s tuning function over visual field locations has stabilized. The output, referred to as the codebook, is thus a matrix with dimensionality *m x d*, where *m* is the number of map units and *d* is the number of discrete locations in the visual field.

There are several ways to probe how this simulated cortex is organized. The first method is to visualize each map unit tuning function directly (**Fig. S2**). In this case, the 820-dimensional tuning function has been reshaped so that its spatial tuning is easy to inspect. The second method summarizes the tuning of each map unit more compactly, following the same methods that are used to visualize retinotopy in biological cortex. Specifically, for each map unit, a 2-dimensional Gaussian was fit to the tuning curve, so that the organization could be summarized by the parameter fits for the angle and eccentricity of the center of the Gaussian as well as its width. This is the method used for the maps shown in Fig. 1 and Fig. 2, as well as Fig. S3-S6, S8-10, S12.

## Supplementary Text

### Parameter Variations

#### Map Shape

What is the impact of changing the map shape, keeping the way the visual field is discretized and sampled? The results of this parameter exploration are shown in **Fig. S3**. The major organizational motifs are present across all map shapes (fovea-periphery, upper/lower, mirrored alternations, smallest scale in the middle surrounded by larger receptive fields). The organization evident in the elliptical map shape is a little more fractured.

#### Map Size

What is the impact of changing the map size, keeping the way the visual field is discretized and sampled? That is, what happens if more or less cortical real-estate is specified to encode the input feature space? **Fig. S4** shows the results of this parameter sweep. The number of map units has a clear impact on the resulting organization. With relatively fewer map units, there is only very slight if any mirrored alternations of vertical and horizontal meridian tuning. With relatively more map units, the organization becomes more fractured. These results imply that, to match the known biological organization, there is a sweet spot related to the degree of compression between the input space (*numSamples* x *numLocations*) and the tuned map (*numUnits* x *numLocations*). In this case of the worked main example, the default heuristic was employed, and interestingly naturally manifests a smooth multi-map organization of the tuned cortex.

#### Scale Space

What is the impact of changing what range of scale space is sampled, (keeping the visual field discretized in the same way, and with all the default parameters of map shape, size, and batch training)? **Fig. S5** shows a small exploration of this parameter space.

Specifically, the range of scales present in the input space was varied, indicated as a range (e.g. [2 24]). Here the minimum scale varied from 2 to 8 degrees (across the columns), and the maximum scale varied from 4 to 24 degrees (along the rows), for a total of 9 different scale space variations. Next, at each visual field location we drew 8 different input samples from the scale range (density d = 8), so that each of the 9 scale space variations had the same number of input samples in the feature space. This yielded 9 different input spaces, which reflect different ways of specifying a multi-scale filter bank over common discretized visual field locations. Next, self-organizing maps were trained for each of these input feature spaces, with either the default number of map units (left panel), or with maps that had twice the cortical real-estate (right panel). For each of the trained maps, the map unit tuning was estimated by fitting 2-d Gaussians. The polar angle parameter is visualized for all 18 maps.

These variations highlight that the way that the multi-scale filter bank is specified is critical for the resulting topography (e.g. the number of alternations), and also depends on the number of units allocated to map the input space. For example, considering scale space sampled from [4 to 24], the smaller map on the left has two meridian reversals, while the same input space mapped onto larger cortical territory (right) has three reversals. Thus, the exact number of reversals and the relative size of each “area” depends on how the multi-scale filter bank is specified and on how much cortical real-estate is allotted to map this input. However, it is also worth noting that all of these map variations have at least some of the major organization motifs present, e.g. upper lower visual field separation, some number of alternations, etc.

#### Single scale example

For illustrative purposes, a simpler model is shown for an input space with one tiling of the visual field at a fixed scale (**Fig. S12**). In particular, this example shows the input space is a 2-dimensional manifold embedded in the high-dimensional input space (**Fig. S12c**), and the resulting 2-dimensional map organization which generally finds this data manifold (**Fig. S12d**).

**Figure S1.**
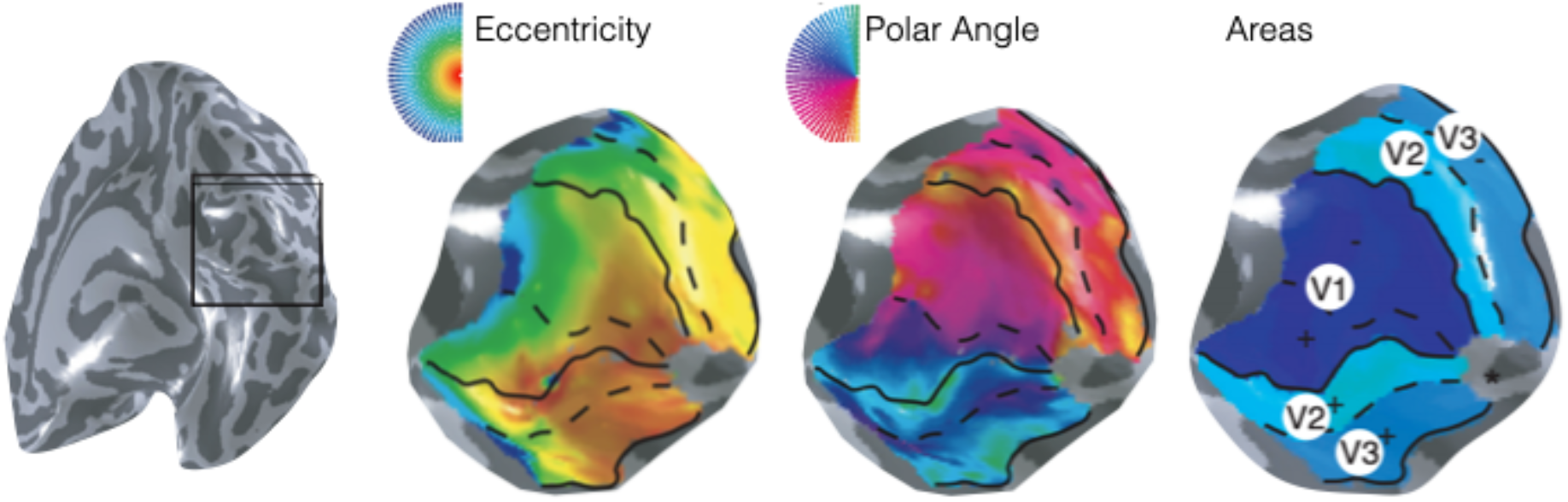
Human visual areal organization. Figure adapted from Wandell et al., 2009.

**Figure S2.**
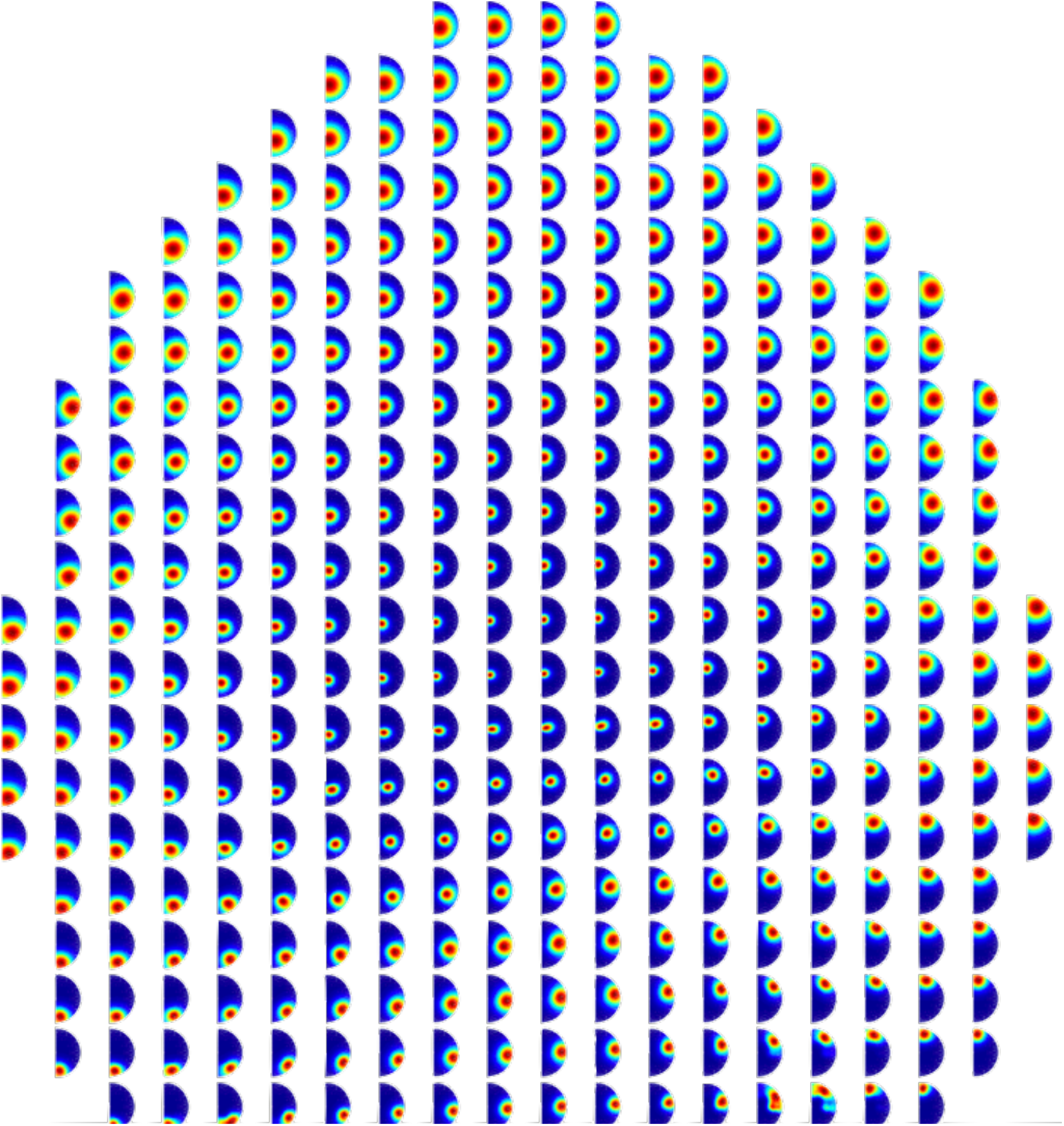
Unit Tuning. Visualization of each map unit’s tuning after training.

**Figure S3.**
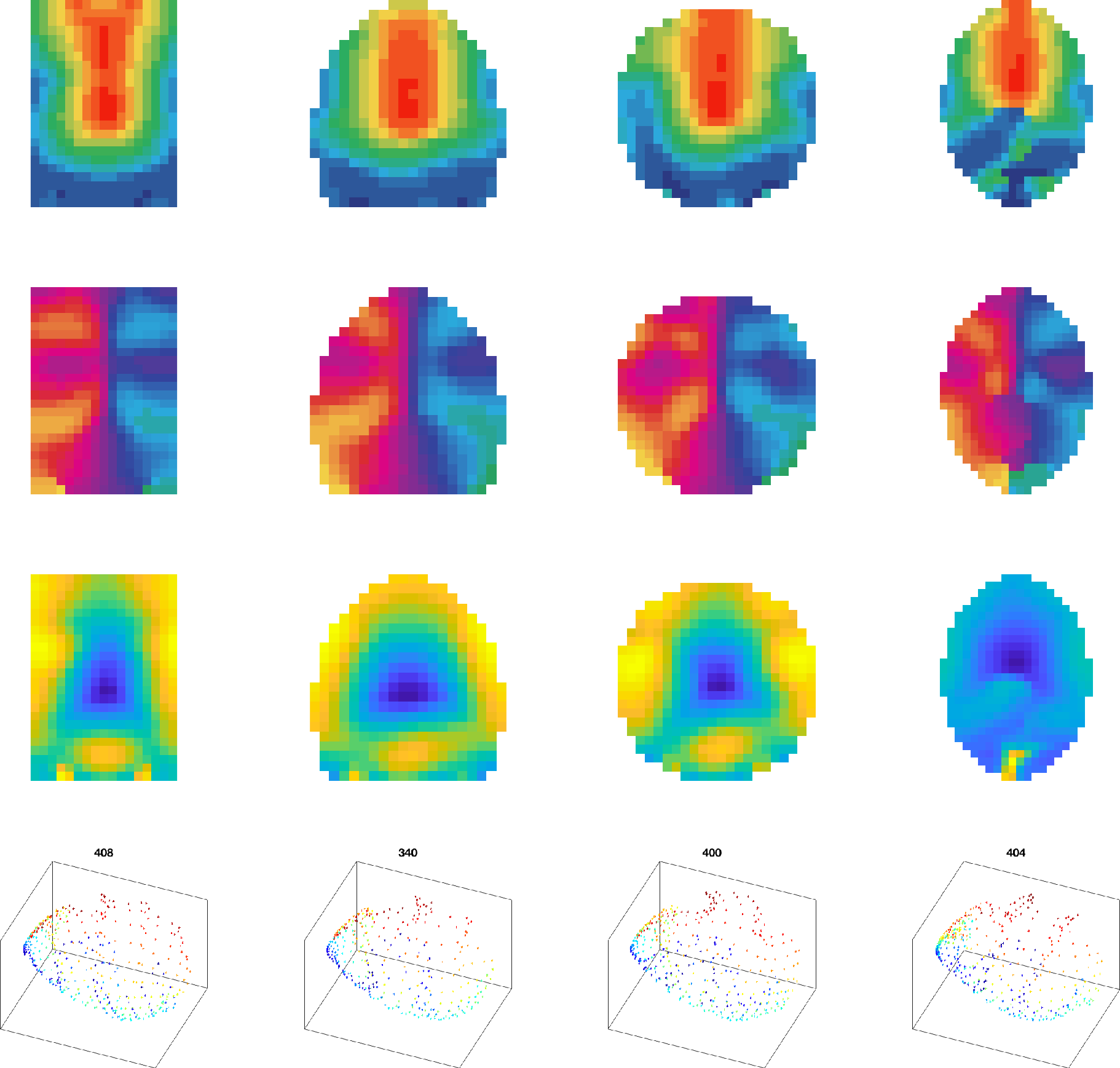
Map shape variations. Each column is a trained map. The shape of the map varies from left to right. Top row: eccentricity fits. Second row: Polar angle fits. Third row: scale fits. Bottom row: visualization of the map units projected into the first three principal components of the input space. Title indicates number of map units.

**Figure S4.**
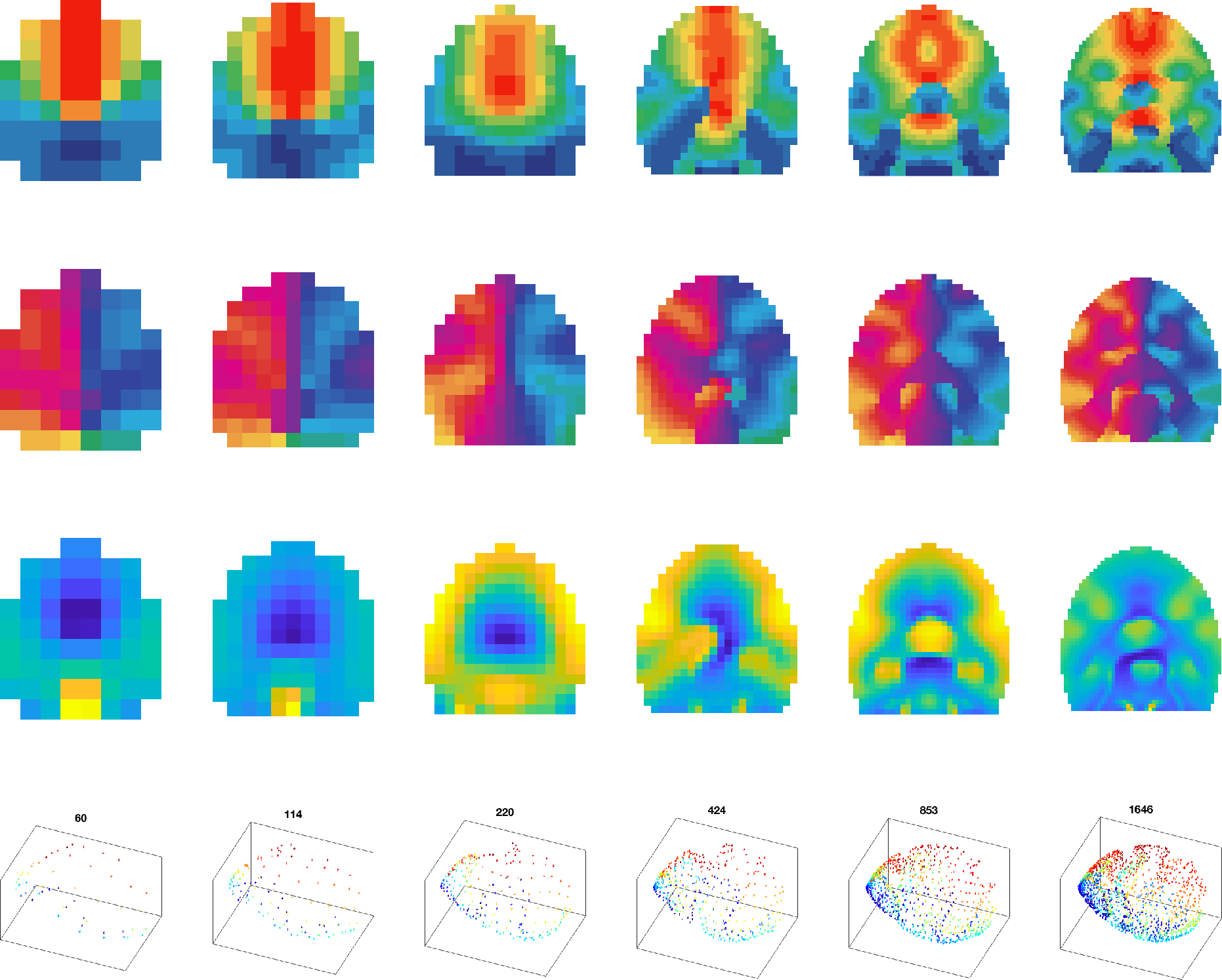
Map size variations. Each column is a trained map. The size of the map varies from small to large, from left to right. Top row: eccentricity fits. Second row: Polar angle fits. Third row: scale fits. Bottom row: visualization of the map units projected into the first three principal components of the input space with number of map units indicated.

**Figure S5.**
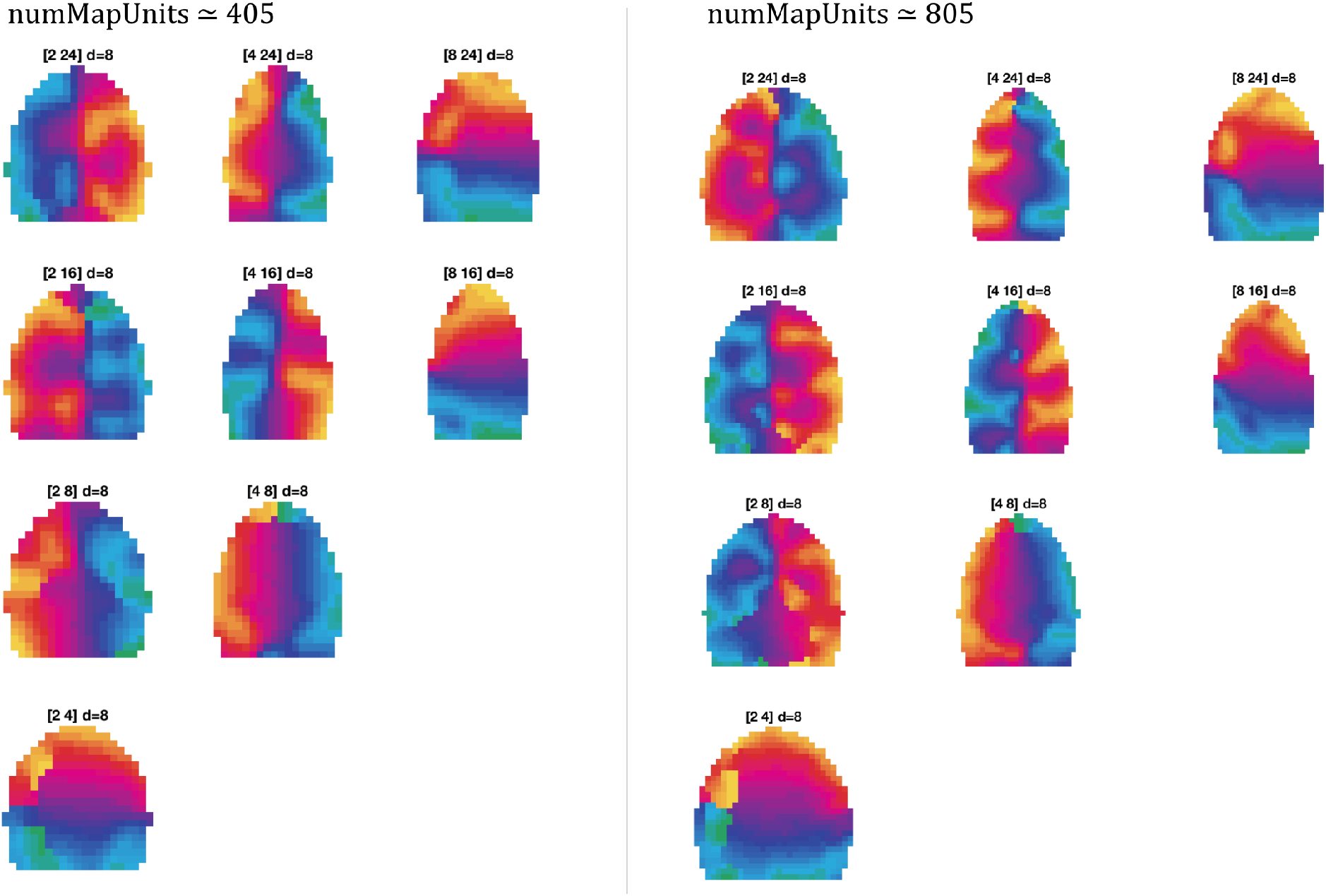
Scale Space and Map Size Variation. Left and right panels indicate two map sizes (left=default size, right=doubled size). Within each panel, the range of scale space sampled is indicated [minScale maxScale], with d=8 samples drawn uniformly from this range, at each location. Map units are colored by the polar angle of the estimated center of their receptive field.

**Figure S6.**
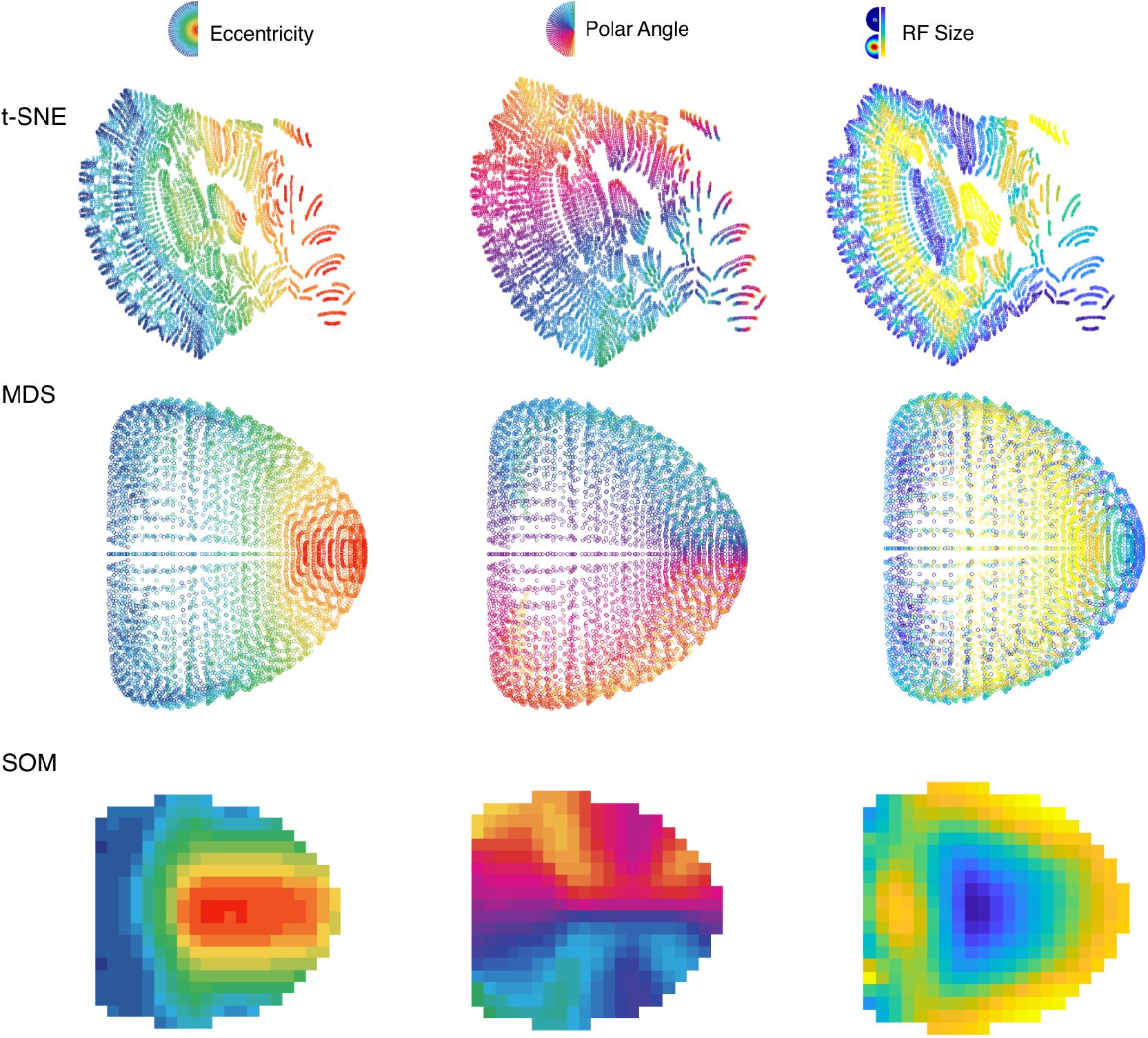
Comparison of dimensionality reduction methods. The same multi-scale filter bank from the main manuscript is plotted with three different dimensionality reduction methods (rows). For the tSNE and MDS row, each dot is an input sample, colored based on its eccentricity, polar angle, and spatial scale (left, middle, right columns). The bottom SOM row replots data from the main manuscript for reference.

**Figure S7.**
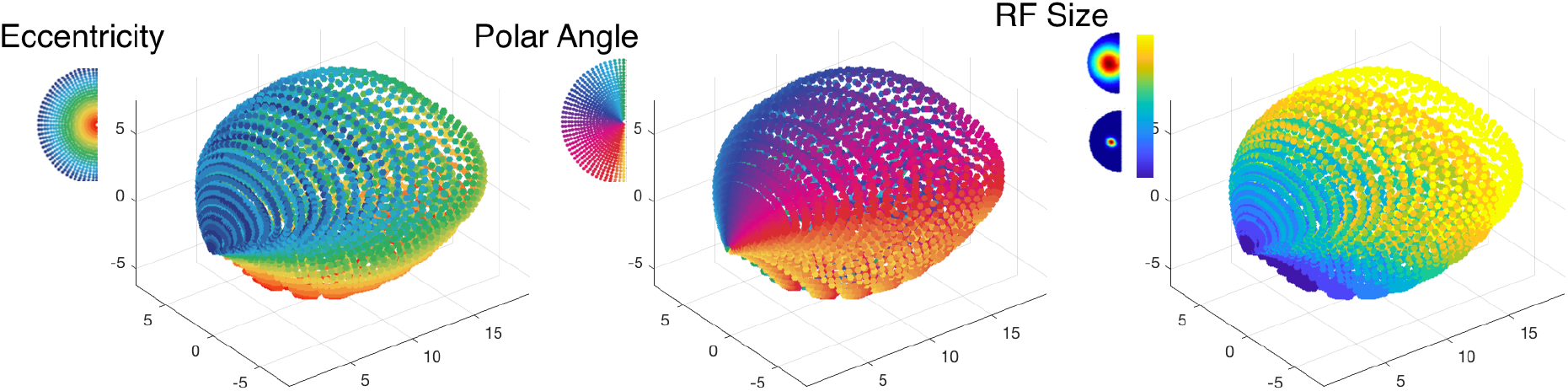
PCA visualization of the input space. The multi-scale filter bank is visualized along its first three principal components. Each point in the plot is an input sample. In each plot, the samples are colored by their eccentricity, polar angle, and scale.

**Figure S8.**
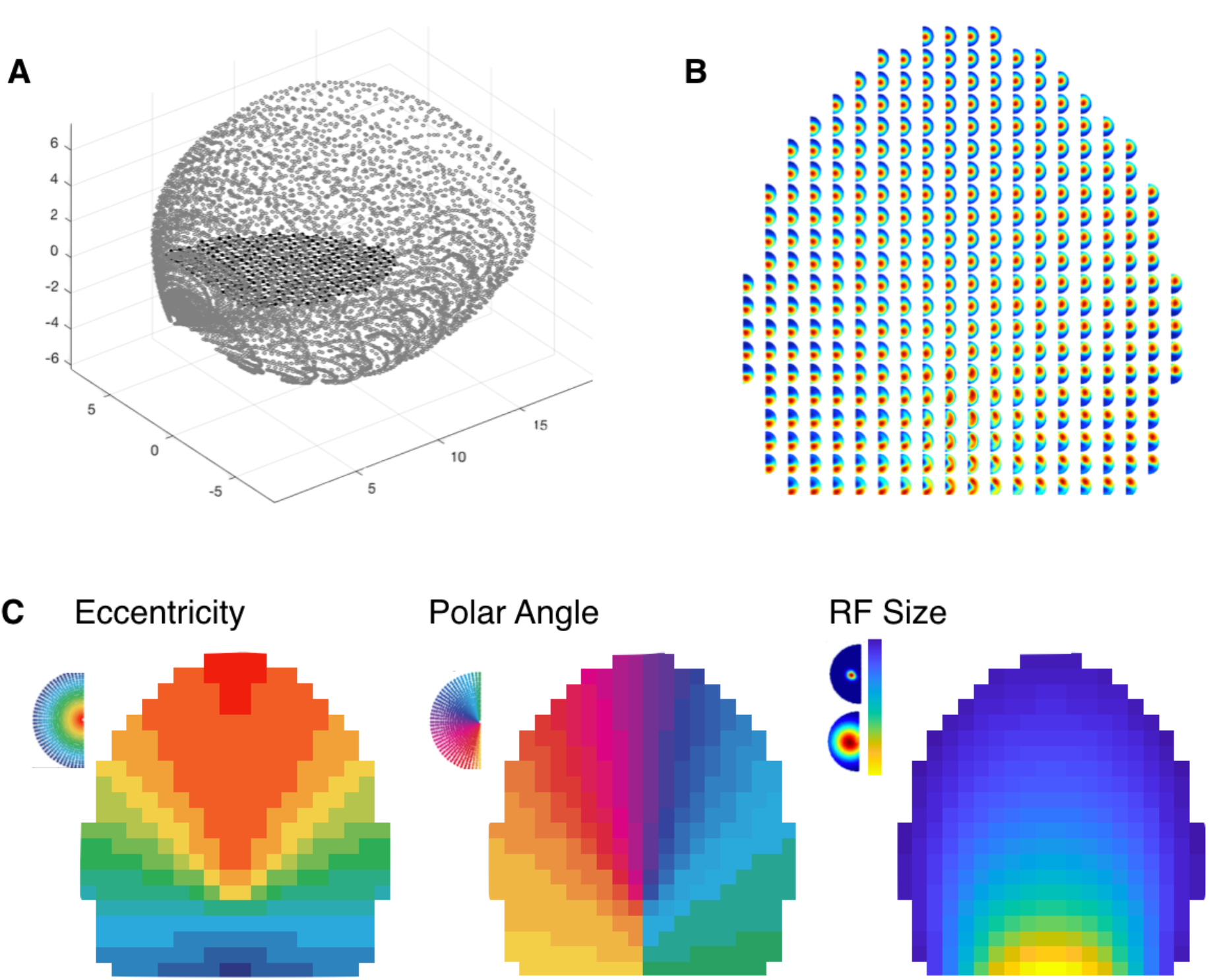
Initialization visualization. (A). Plot of the map units’ tuning in the context of the input feature PC space. The x, y, and z axes reflect the first three components of the input feature matrix. Gray dots are samples of the input feature space. Black dots correspond to a map unit and are connected in a grid to their nearest neighbors. (B). Visualization of each map unit’s initial tuning. Each subplot corresponds to a map unit, and the weights along the 820 visual field locations have been arranged to visualize the visual field tuning of each unit. (C). After fitting 2-d Gaussians to each map unit tuning, the eccentricity and polar angle of the center of the Gaussian is plotted (left, middle), along with the fitted sigma/scale (right).

**Figure S9.**
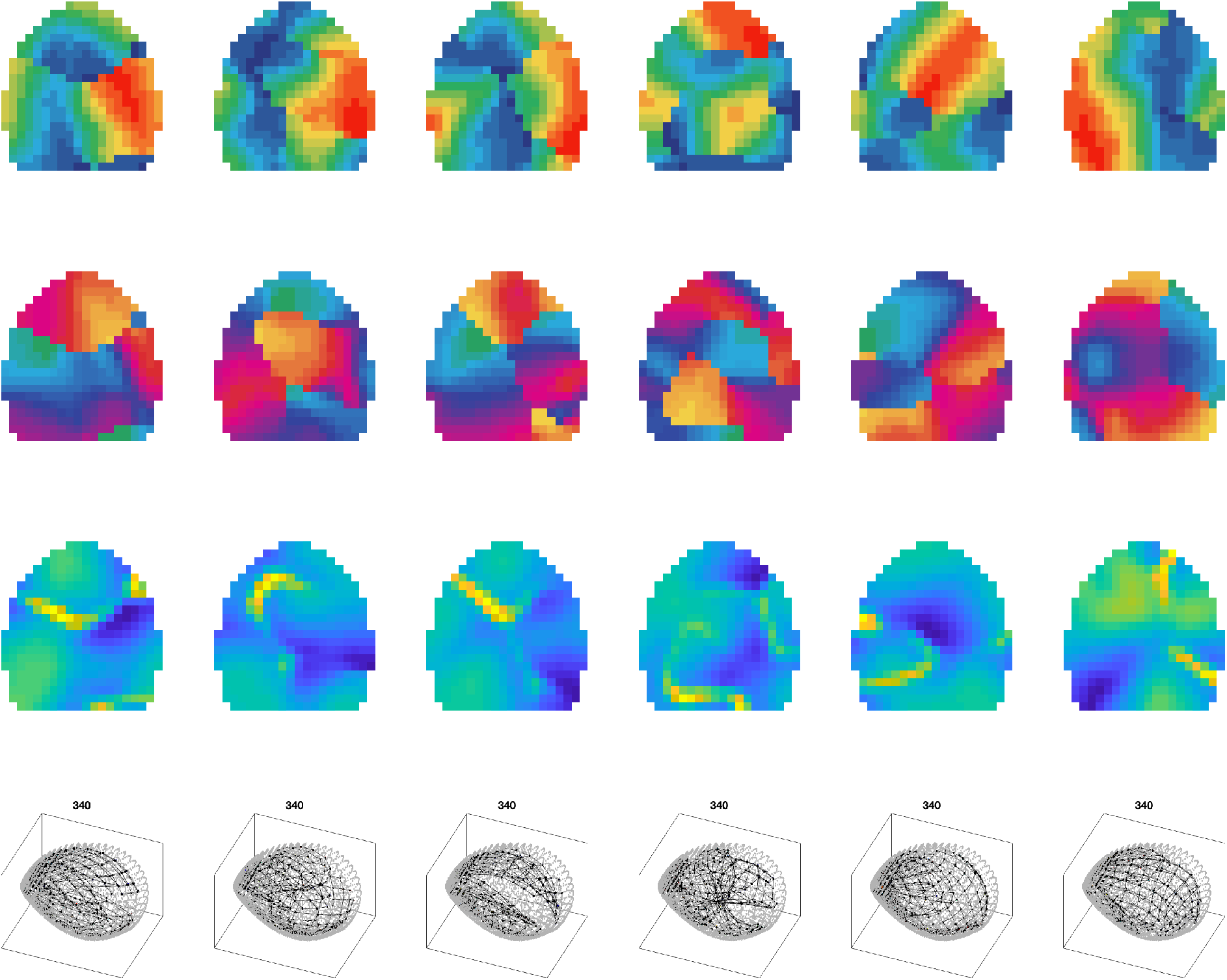
Random Map Initialization. Each column is a trained map with a different random initialization. Top Row: Eccentricity fits. Second row: Polar angle fits. Third row: scale fits. Bottom row: visualization of the map units projected into the first three principal components of the input space, with number of map units indicated.

**Figure S10.**
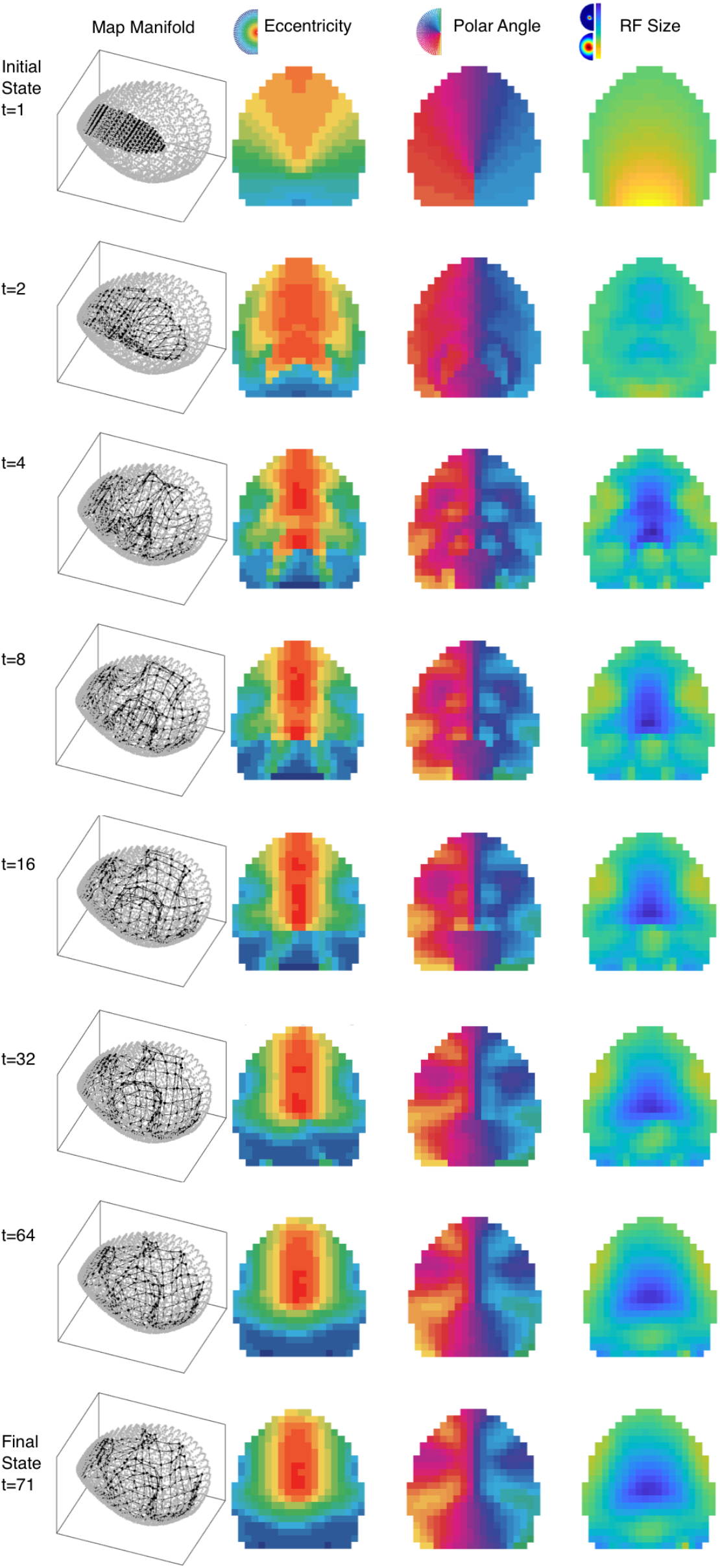
Map development visualization. Rows: training iterations. Columns: fitted 2D Gaussian map unit parameters

**Figure S11.**
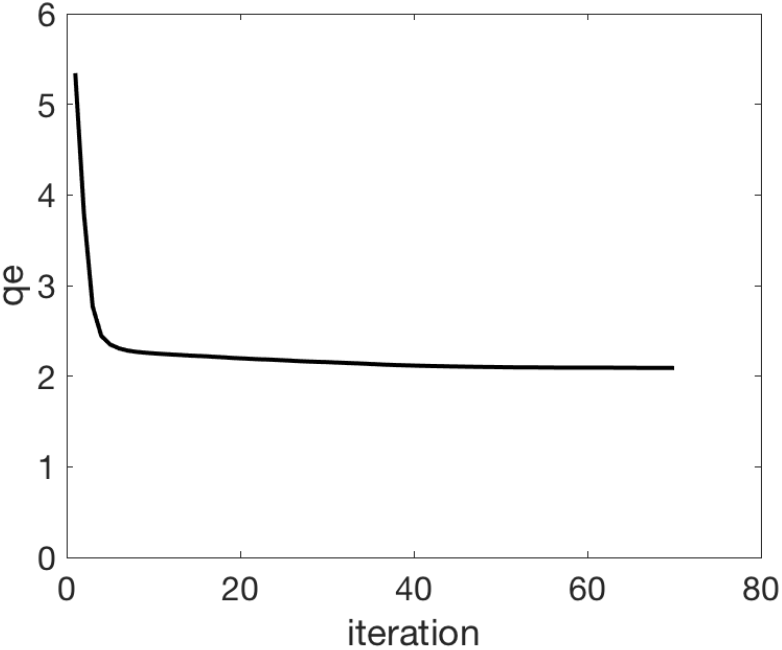
Quality of the estimation (qe) as a function of training iteration

**Figure S12.**
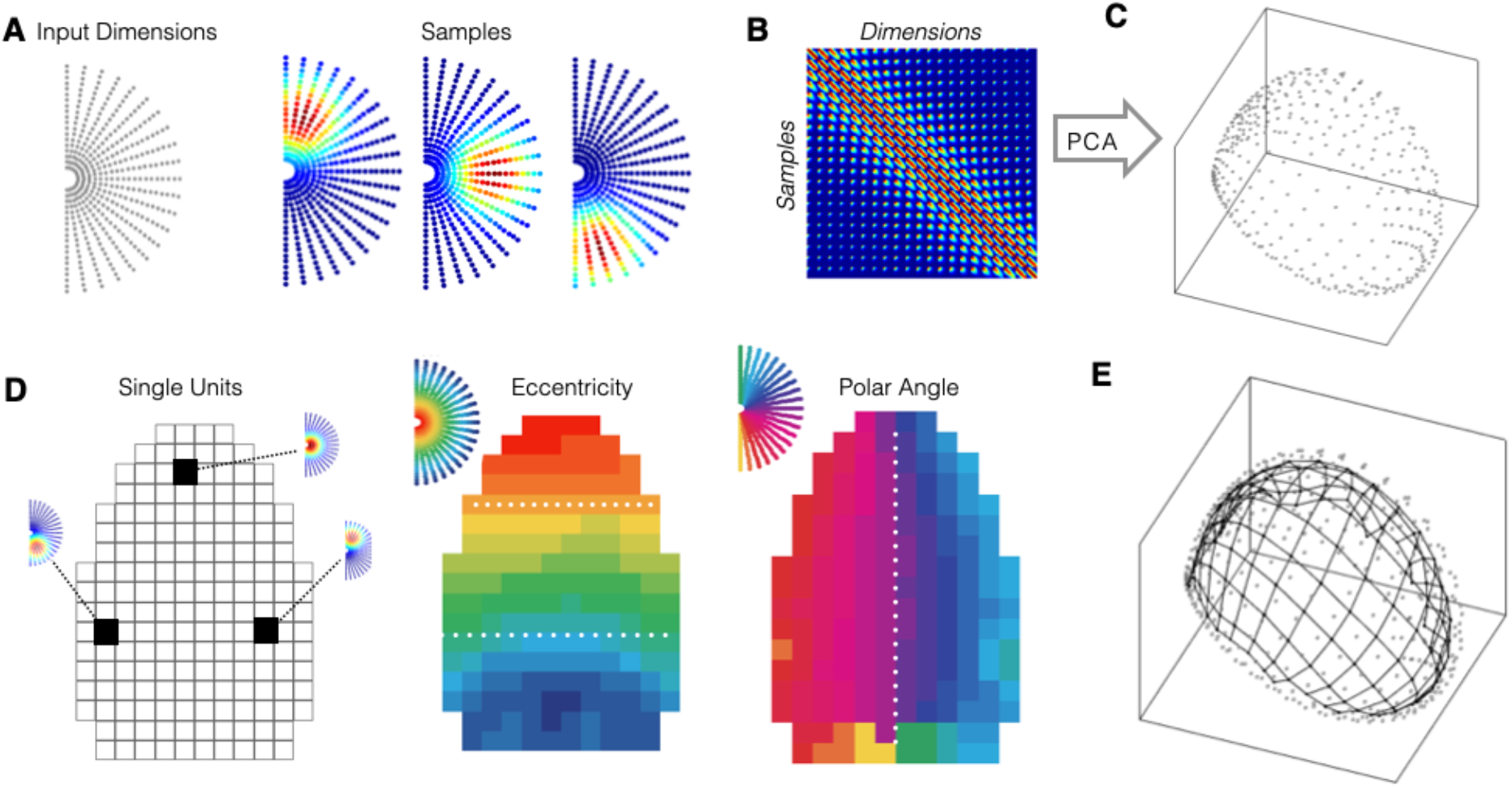
Single scale example. (A) The visual field was discretized to establish the input dimensions, and a set of 2-dimensional Gaussian input samples were created, centered at each location, with a fixed size. Three examples are shown. (B). Visualization of this input space, as a matrix of *numSamples* x *numDimensions*. (C). Projection of this input space along its first three principal components. (D). After fitting an SOM, each unit has tuning weights along the input dimensions, with 3 example learned tuning weights shown. These are summarized in the adjacent plots by both the eccentricity and polar angle coordinates of the fitted 2d-Gaussian models, revealing a single visual area. (E). The data space is shown (as in C) with light gray dots, with the SOM units are also plotted based on their tuning with black dots, connected to adjacent map units in the SOM with black lines.

### Movie S1

An animated gif of the map development, visualizing the state of the map from initialization through training iterations can be found at http://konklab.fas.harvard.edu/misc/Dev.gif.

### Code Base

RetinoSOM toolbox.

Copyright (C) 2020 Talia Konkle.

This program is free software: you can redistribute it and/or modify it under the terms of the GNU General Public License as published by the Free Software Foundation, either version 3 of the License, or (at your option) any later version. This program is distributed in the hope that it will be useful, but WITHOUT ANY WARRANTY; without even the implied warranty of MERCHANTABILITY or FITNESS FOR A PARTICULAR PURPOSE. See the GNU General Public License for more details. See <https://www.gnu.org/licenses/>.

